# HTSSIP: an R package for analysis of high throughput sequencing data from nucleic acid stable isotope probing (SIP) experiments

**DOI:** 10.1101/166009

**Authors:** Nicholas D. Youngblut, Samuel E. Barnett, Daniel H. Buckley

**Author notes:** Corresponding author: Nicholas D. Youngblut, Max Planck Institute for Developmental Biology, Spemannstraße 35, 72076 Tübingen, Germany.

## Abstract

Combining high throughput sequencing with stable isotope probing (HTS-SIP) is a powerful method for mapping *in situ* metabolic processes to thousands of microbial taxa. However, accurately mapping metabolic processes to taxa is complex and challenging. Multiple HTS-SIP data analysis methods have been developed, including high-resolution stable isotope probing (HR-SIP), multi-window high-resolution stable isotope probing (MW-HR-SIP), quantitative stable isotope probing (q-SIP), and ΔBD. Currently, the computational tools to perform these analyses are either not publicly available or lack documentation, testing, and developer support. To address this shortfall, we have developed the *HTSSIP* R package, a toolset for conducting HTS-SIP analyses in a straightforward and easily reproducible manner. The *HTSSIP* package, along with full documentation and examples, is available from CRAN at https://cran.r-project.org/web/packages/HTSSIP/index.html and Github at https://github.com/nick-youngblut/HTSSIP.

## Introduction

Stable isotope probing of nucleic acids (DNA- and RNA-SIP) is a powerful method for mapping *in situ* metabolic processes, such as nitrogen and carbon cycling, to microbial taxa. Historically the sensitivity of nucleic acid SIP has been limited by the low throughput of DNA sequencing and the low taxonomic resolution of DNA fingerprinting techniques [1,2]. Recently, DNA- and RNA-SIP have been combined with high throughput sequencing of PCR amplicons (HTS-SIP), which allows researchers to map *in situ* metabolic processes to thousands of taxa resolved at a fine taxonomic resolution [3–5].

While HTS-SIP is proving to be a very useful method for exploring *in situ* metabolic processes in complex microbial communities, the accurate analysis of HTS-SIP datasets is complex [6,7]. Multiple strategies have been developed for analyzing HTS-SIP data, including high-resolution stable isotope probing (HR-SIP) [5], multi-window high-resolution stable isotope probing (MW-HR-SIP) [7], quantitative stable isotope probing (q-SIP) [3], and ΔBD [5]. The goals of these methods differ, with HR-SIP and MW-HR-SIP designed to accurately identify taxa that have incorporated isotopically labeled substrate (*i.e*. ‘incorporators’), while the main goal of q-SIP and ΔBD is to quantify the amount of isotopic enrichment for each taxon (*i.e*. atom % excess). While all methods use amplicon sequences (*e.g*. 16S rRNA or fungal ITS sequences) from multiple fractions of each isopycnic gradient, HR-SIP, MW-HR-SIP, and ΔBD solely use sequence data while q-SIP additionally requires qPCR derived estimations of gene copy number from each gradient fraction. Recently, Youngblut and Buckley developed a HTS-SIP simulation model and showed that MW-HR-SIP is more accurate for identifying incorporators than HR-SIP and q-SIP, while q-SIP is generally more precise than ΔBD for quantifying isotopic enrichment [7].

The code for performing each of these HTS-SIP analyses is limited in availability, documentation, and developer support; all of which severely limit the ease of use and reproducibility of HTS-SIP analyses. To address this deficiency, we developed the *HTSSIP* R package, which includes the following features:
- Functions for conducting HR-SIP, MW-HR-SIP, q-SIP, and ΔBD to analyze data from DNA-SIP and RNA-SIP experiments
- Functions for performing HTS-SIP dataset simulation, as described [7]
- Functions for exploratory analysis of simulated HTS-SIP data, useful for predicting how different experimental designs can alter experimental outcomes
- Functions for exploratory analysis of real HTS-SIP data, useful for conducting post-hoc analyses
- Ability to run analyses with parallel processing
- Extensive documentation and tutorials (see the *HTSSIP* vignettes)

### Package description

#### Input data

Dataset input is handled by the *Phyloseq* R package, a feature-rich package for general microbiome data analysis that can be used to import many common microbiome data formats [8]. *HTSSIP* includes convenience functions to easily and flexibly designate the experimental design of the SIP experiment for downstream HTS-SIP analyses (Figure 1).

**Figure 1.**
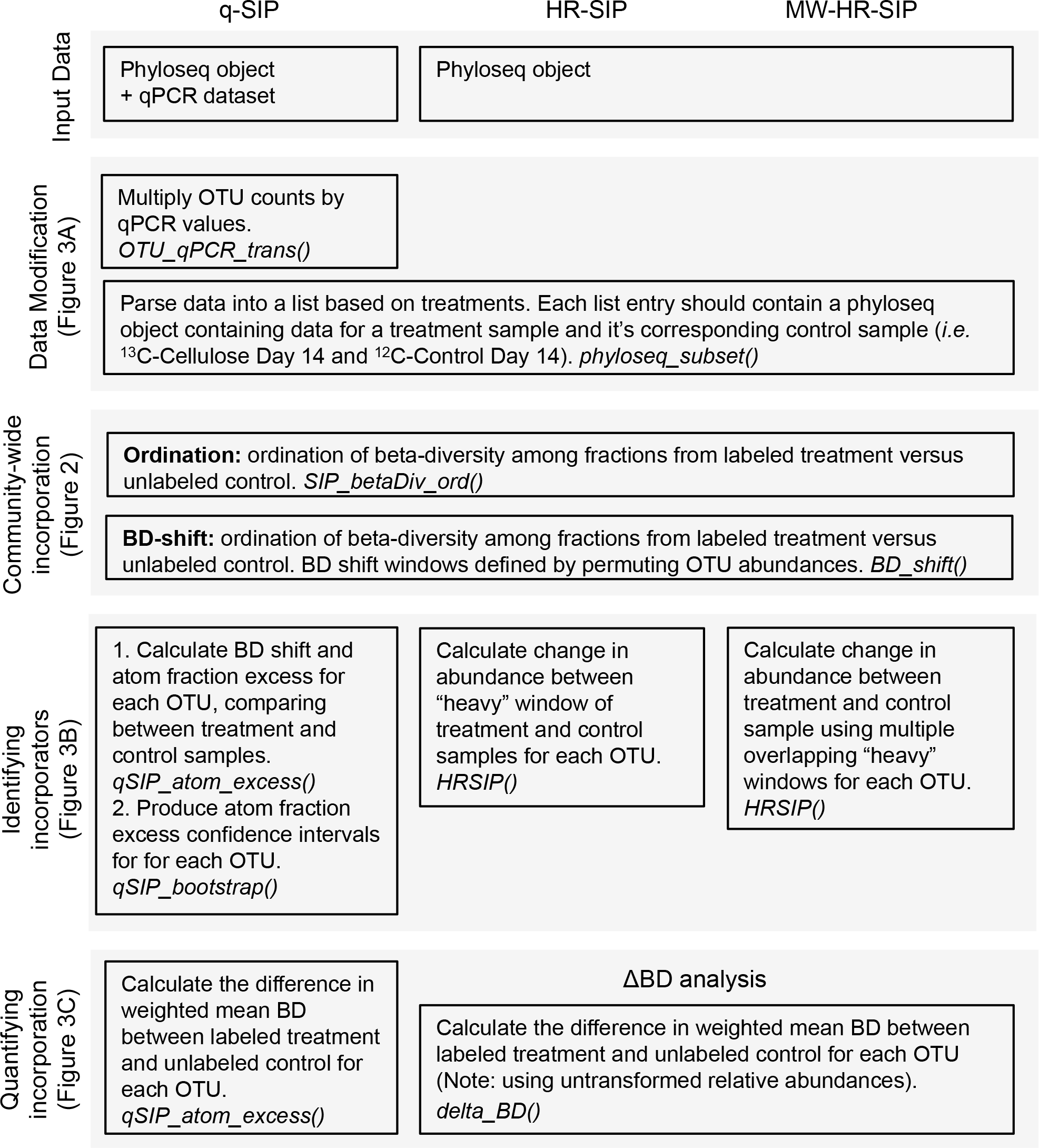
*A diagram depicting the possible analyses available in the* HTSSIP *R package*. The R functions to conduct each workflow step are italicized, and the figure references refer to example data produced by these workflow steps.

#### HTS-SIP dataset exploratory analyses

A common first step in analyzing nucleic acid SIP data is to quantify the total nucleic acid concentration or gene copy number (estimate by qPCR) across density gradients in order to determine the buoyant density (BD) “shift” of nucleic acids in isotopically labeled treatments versus unlabeled controls [9,10]. The general expectation is that a “shift” of nucleic acid BD from “light” towards “heavy” densities is indicative of isotope incorporation. However, in a well designed SIP experiment, the ratio of exogenous to indigenous substrate should be small, and this can produce an imperceptible BD shift [4]. In addition, an extensive shift may indicate excessive cross-feeding [11]. HTS-SIP methods can detect taxa that have incorporated low levels of isotope, or occur at frequencies that are so low that they do not cause a shift in the overall BD of community nucleic acids [5]. As a result, analysis of the BD distribution of total nucleic acids within density gradients is of little utility in assessing the results of nucleic acid SIP experiments performed on complex communities.

As a simpler alternative, which leverages the power of high-throughput sequencing techniques, BD “shifts” can be inferred solely from sequence data [4,5]. Given that incorporators will be more abundant in “heavy” gradient fractions of the labeled treatment versus the unlabeled control, a BD shift can be inferred by assessing the beta-diversity between treatment and control gradient fractions. This approach is more sensitive for detecting community-level isotope incorporation than the approach of quantifying total nucleic acid concentration across the density gradient [7]. *HTSSIP* implements two methods for using beta-diversity to assess isotope incorporation at the community-level: an ordination approach and an approach that expresses beta-diversity between corresponding treatment and control fractions as a function of their BD (Figure 1).

The ordination approach simply involves pairwise calculations of a beta-diversity metric between all gradient fractions from isotopically labeled treatments and corresponding unlabeled controls, followed by visualizing the distance matrix with either principal coordinates analysis (PCoA) or non-metric multidimensional scaling (NMDS). An increase in beta-diversity between corresponding gradient fractions of labeled samples and controls is expected if isotope incorporation causes a change in the BD of OTUs (Figure 2A & 2B).

**Figure 2.**
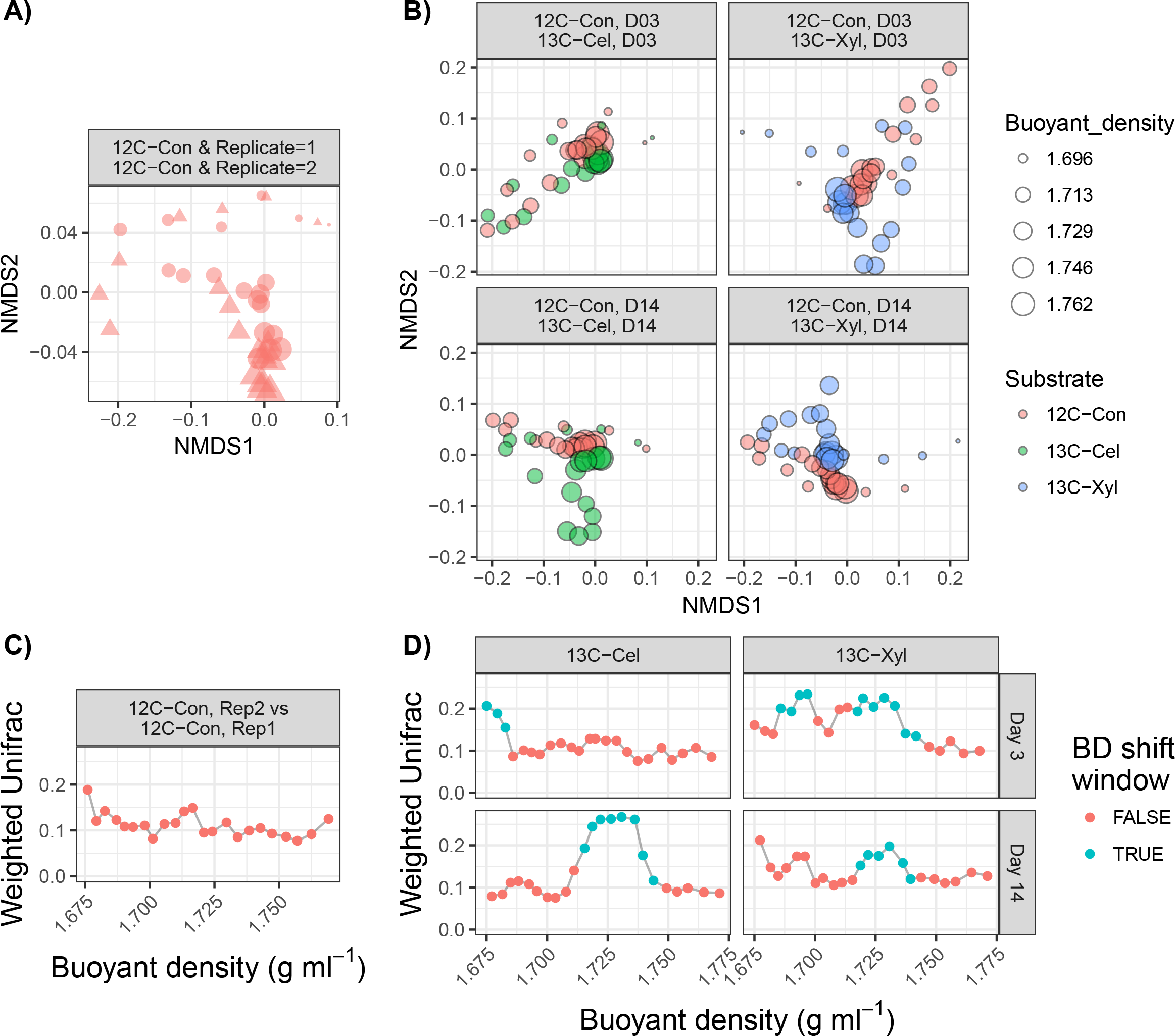
*Examples of the ordination and BD-shift analyses for assessing community-level incorporation*. Plots A and B are non-metric multidimensional scaling (NMDS) ordinations of beta-diversity (16S rRNA OTUs; 97% sequence identity; weighted Unifrac) calculated between gradient fractions from a HTS-DNA-SIP experiment conducted with agricultural soil. Plot A compares fractions from replicate unlabeled control gradients, with different symbols (circles and triangles) used to distinguish different replicates, and with symbol diameter scaled in relation to fraction buoyant density as indicated in the accompanying scale. Plot B compares fractions from labeled treatments (“13C-Cel” for ^13^C-cellulose or “13C-Xyl” for ^13^C-xylose) versus their corresponding unlabeled controls (“12C-Con”) at 3 or 14 days after substrate addition (“D03” and “D14”, respectively). The NMDS stress values ranged from 0.06 to 0.07. An increase in beta-diversity is expected between labeled and unlabeled “heavy” fractions in response to isotope incorporation. Plots C and D depict the same data as in Plots A and B, but the beta-diversity comparisons between labeled treatment and unlabeled control are indicated only for fractions that correspond in BD. To account for partial overlap between labeled and unlabeled fractions, the weighted mean beta-diversity value is calculated based on percent overlap in BD ranges. "BD shift windows" indicate regions defined by ≥3 consecutive fractions with significantly high beta-diversity resulting from isotope incorporation, with significance defined by permuting OUT abundances and recalculating beta-diversity values (100 bootstrap replicates; P < 0.05). The dataset used is a subset from the dataset from Youngblut and Buckley [7].

While the ordination approach provides a useful overview of community-wide isotope incorporation, the extent of incorporation is difficult to compare among multiple treatments (*e.g.* ^13^C-cellulose versus ^13^C-xylose). The second approach implemented in *HTSSIP* visualizes DNA BD shifts by calculating pairwise beta-diversity of corresponding gradient fractions between treatment and control gradients. To deal with partially overlapping gradient fractions between gradients, the weighted mean beta-diversity is calculated from all treatment gradient fractions that overlap each control gradient fraction, with weights defined as the percent overlap in the BD range of each fraction (Figure 2C & 2D). A permutation test is used to identify BD ranges of high beta-diversity resulting from BD shifts (“BD shift windows”). The permutation test involves constructing bootstrap confidence intervals (CI) of beta-diversity by permuting OTU abundances among labeled treatments (*i.e*. a null model where OTUs in treatment are randomly dispersed relative to the control). A note in interpreting these data is that isotope incorporation will cause DNA to shift out of “light” gradient fractions and into “heavy” gradient fractions. Hence, in the presence of isotope incorporation, high beta-diversity can be observed in both “heavy” and “light” gradient fractions. Alternatively, in the absence of isotope incorporation, beta-diversity will remain low across all gradient fractions.

#### Identifying incorporators

HR-SIP, MW-HR-SIP, and q-SIP can all be used to identify incorporators. To illustrate the application of HR-SIP, MW-HR-SIP, and q-SIP in the *HTSSIP* R package, we simulated a simplified HTS-SIP dataset consisting of 10 OTUs (Figure 3A). Our purpose here is merely to illustrate functions of the *HTSSIP* R package; comprehensive assessment of the accuracy of these techniques is available elsewhere [7]. Briefly, HR-SIP identifies incorporators by utilizing DESeq2 to identify OTUs that have high differential relative abundance in “heavy” fractions of labeled treatment versus unlabeled control [12]. MW-HR-SIP takes the same relative abundance based approach as HR-SIP but uses multiple overlapping “heavy” BD windows (while correcting for multiple hypotheses). In contrast, q-SIP uses qPCR data to transform OTU relative abundance distributions into pseudo-absolute abundance distributions (Figure 3A), and then BD shifts are determined from these transformed distributions by calculating the difference in center of mass for each OTU in treatment versus control gradients. Atom fraction excess can thus be calculated for specific isotopes (*e.g*. ^13^C or ^15^N) based on the calculations described in the work of Hungate and colleagues [3]. In order to identify incorporators, a permutation test is used to construct bootstrap confidence intervals of atom fraction excess. Sensitivity in identifying incorporators can depend on the methods used (Figure 3B; and see [7]). SIP experiments can be simulated using the SIPSim toolset [7], and these data analyzed using the *HTS-SIP* R package. Such *in silico* evaluation is valuable for predicting possible experimental outcomes and the expected analytical accuracy of SIP experiments based on details of experimental design prior to conducting experiments.

**Figure 3.**
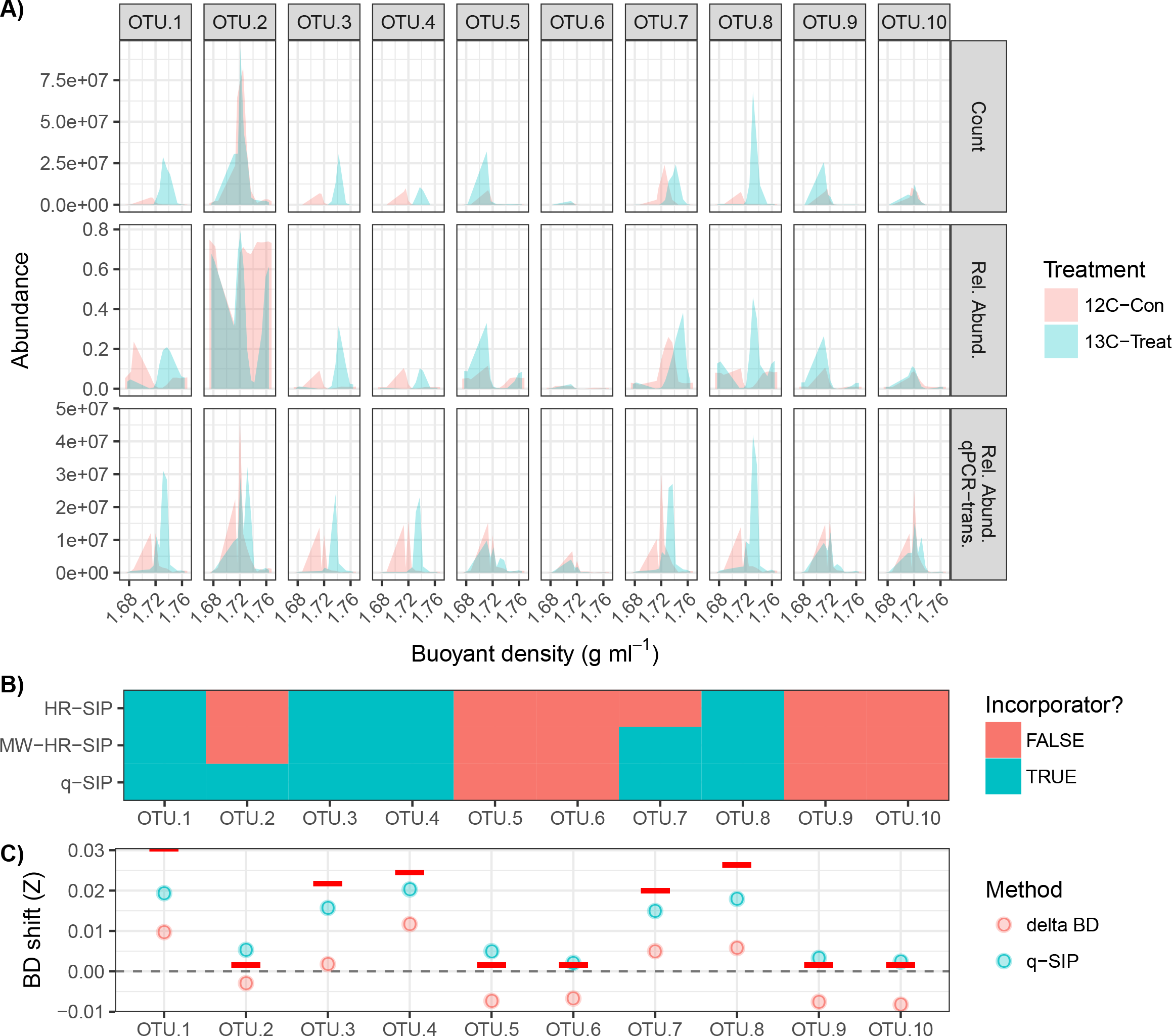
*Examples of using the HTSSIP R package for data processing, data exploration, incorporator identification, and quantification of BD shifts (Z)*. The SIPSim toolset was used to simulate the relative abundances of 10 OTUs across 24 gradient fractions in an experiment that includes a single ^13^C-treatment (“13C-Treat”) and a single ^12^C-control (“12C-Con”) condition, each with 3 experimental replicates. Half of the OTUs had an atom fraction excess of 30 to 100%, while the others were 0%. qPCR estimates of total community 16S rRNA copy numbers were also simulated with SIPSim, and qPCR analytical error was modeled based on error estimated from Hungate and colleagues [3]. Plot A depicts the raw abundances (“Counts”), fractional relative abundance (“Rel. Abund.”), and relative abundances transformed by simulated qPCR data (“Rel. Abund. qPCR-trans.”). For clarity, only 1 of the 3 experimental replicates is shown. Plot B shows which OTUs were identified as incorporators by the statistical methods described for HR-SIP, MW-HR-SIP, or q-SIP. A Benjamini-Hochberg corrected p-value cutoff of 0.1 was used for HR-SIP and MW-HR-SIP, and 100 bootstrap replicates were used to calculate confidence intervals for q-SIP, as described [3]. Plot C shows the mean BD shift of each OTU as quantified by ΔBD or q-SIP. The dashed line signifies a BD shift (Z) of 0.0 g ml^-1^, and the red bars show the true theoretical BD shift resulting from ^13^C isotope incorporation.

*HTSSIP* implements HR-SIP based on the code provided in the work of Pepe-Ranney and colleagues [5]. MW-HR-SIP is implemented in *HTSSIP* based on the R code provided in the SIPSim HTS-SIP dataset simulation toolset [7]. The *HTSSIP* implementation of q-SIP is based on the method’s description in the work of Hungate and colleagues [3]. Implementations of each method include the option for parallel processing of each algorithm. Parallelization is implemented through the *plyr* R package [13], which allows for various parallel backends to be used such as *doSNOW* and *doParallel*.

#### Quantifying isotopic enrichment

Unlike HR-SIP and MW-HR-SIP, the main goal of q-SIP and ΔBD is to quantify isotopic enrichment. To illustrate the use of *HTSSIP* for conducting q-SIP and ΔBD, we applied both analyses to the simplified HTS-SIP dataset described above (Figure 3C). ΔBD is implemented in *HTSSIP* as described in the work of Pepe-Ranney and colleagues [5]. As shown in Youngblut and Buckley [7], q-SIP and ΔBD can produce substantially different estimates of isotope incorporation.

#### Simulating datasets

*HTSSIP* provides functions to simulate simple HTS-SIP datasets for use in software testing, analysis pipeline development, and gaining familiarity with software and data formats. However, the SIPSim toolset is recommended for evaluating possible SIP experimental designs and for testing the accuracy of HTS-SIP analyses, because the simulation framework for SIPSim is based on the physics of isopycnic centrifugation, unlike the simulations possible with *HTSSIP* [7]. *HTSSIP* utilizes *coenocliner*, an R package designed for simulating taxon abundance across environmental gradients, to simulate taxon abundance distributions across buoyant density gradient fractions [13].

#### Availability

The *HTSSIP* package and the data used in this work are available from CRAN at https://cran.r-project.org/web/packages/HTSSIP/index.html and Github at https://github.com/nick-youngblut/HTSSIP.

### Future work

Future development of the *HTSSIP* package will include i) functions for mapping incorporator status to phylogenies and visualizing the results ii) direct integration with the SIPSim toolset for rapid HTS-SIP experimental design and assessment of accuracy iii) functions analyzing shotgun metagenome data derived from SIP experiments.

## Conclusions

Given the power of HTS-SIP for mapping *in situ* metabolism to taxonomic identity, adoption of the technique by researchers will greatly help to resolve connections between microbial ecology and taxonomy. Currently, HTS-SIP data analysis is complex, with few existing computational tools to aid researchers. The R package *HTSSIP* provides a single, standardized analysis pipeline that facilitates reproducible analyses on HTS-SIP datasets and direct cross-study comparisons. Moreover, *HTSSIP* can be combined with the SIPSim toolset to simulate and evaluate possible DNA-SIP experimental designs.

## Acknowledgements

This work is supported by the Department of Energy Office of Science, Office of Biological & Environmental Research Genomic Science Program under award numbers DE-SC0004486 and DE-SC0010558.

